# The role of eye movements in the process of silicone oil emulsification after vitreoretinal surgery

**DOI:** 10.1101/2024.06.06.597725

**Authors:** Irene Nepita, Camilla Brusati, Libero Liggieri, Francesca Ravera, Mariantonia Ferrara, Alessandro Stocchino, Mario R. Romano, Eva Santini, Rodolfo Repetto

## Abstract

**Background:** Emulsification of silicone oil (SO) is a feared and common complication of SO tamponade as potentially associated with significant risks to ocular health, including elevated intraocular pressure (IOP), glaucoma, corneal and retinal changes. The aim of this study was to investigate the role and interplay of major factors on the formation of SO emulsion, such as eye rotations and albumin, a blood serum protein known to affect interfacial properties.

**Methods:** Experiments were conducted in a realistic model of the vitreous chamber, filled with SO and an aqueous solution containing different concentrations of albumin. The model was subjected to harmonic and saccadic rotations, at body temperature.

**Results:** No emulsions were detected in the absence of endogenous proteins in the aqueous solution. The presence of albumin significantly influenced emulsion formation, acting as a surfactant. Mechanical energy from eye movements was also found to contribute to emulsification, with higher mechanical energy provided to the system leading to smaller droplet sizes. The emulsions formed were stable over extended times.

**Conclusions:** This study highlights the complex interplay of factors influencing SO emulsification in the vitreous chamber. A better understanding of the mechanisms underlying SO emulsification is crucial for developing strategies to mitigate SO emulsion and the related complications.

## 1 Introduction

Pars plana vitrectomy (PPV) is the surgical procedure of choice to treat several vitreoretinal pathologies. Silicone oil (SO) is commonly used as long-term intraocular tamponade for the management of complex diseases, such as complicated retinal detachment.^1^ However, multiple SO-related ocular complications have been reported so far. ^1^ In particular, the formation of intraocular SO emulsion is highly undesirable and appears to have a crucial role in complications affecting nearly all ocular structures, such as acute and chronic changes in intraocular pressure (IOP), optic neuropathy, retinal and corneal alterations, cataract, and extraocular extension. ^1,2^ In addition, SO emulsification appears to be strictly related to intraocular inflammation. ^1^ Indeed, emulsified SO droplets can be phagocytised by macrophages, microglial cells and retinal pigment epithelium (RPE) cells, triggering an inflammatory response, that can, in turn, promote further SO emulsification due to the release of endogenous proteins in the intravitreal aqueous phase, able to modify rheological interfacial properties between aqueous solution and SO. ^3^ In this regard, it has been demonstrated that various blood constituents, such as lymphocytes, plasma, serum, red blood cells, haemoglobin, fibrinogen, fibrin, *γ*-globulins, plasma lipoproteins and purified HDL-apolipoproteins, are bio-molecules able to act as emulsifiers/surfactants for SO when dissolved in aqueous solution, adsorbing at the water-oil interface and modifying its mechanical interfacial properties. ^4–6^ Nepita et al. ^7^ demonstrated that the presence of two serum proteins, albumin and *γ*-globulins, in an aqueous solution had a significant effect on the interface SO-aqueous resulting in emulsions stable on the time scale of months. Besides the presence of surface active species, the formation of stable emulsions is related to the enforcement of a mechanical energy.

Results obtained with albumin suggest that, under post-surgical conditions, even a small amount of mechanical energy could be sufficient to break the interface into small droplets, which, in addition, are stable against coalescence. ^7^ Thus, a major factor influencing SO emulsification is represented by eye movement as shear stresses at the SO-aqueous interface generated during eye rotations (such as saccadic motions and REMs), which play a key role in destabilising the interface. ^8–11^ Fluid dynamics in the vitreous chamber produced by eye rotations has been studied with numerical models, ^12–18^ experimental approaches, ^19–21^ and in-vivo measurements,^22,23^ demonstrating that the flow field induced by eye rotations has a complicated three dimensional structure. Recent experimental studies investigated the motion of SO partially filling a model of the vitreous chamber, ^8,9^ the shape of the interface SO-aqueous solution in the eye, ^24^ the dynamics induced by eye rotations ^25^ and the mechanisms relating shear flow generated by eye rotations with the breakdown of the SO-aqueous interface, ^26^ demonstrating the need of understanding the behavior of vitreous substitutes and the shear force as a factor influencing emulsification. Finally, Wang et al. ^10,11^ performed experiments in a spherical model of the eye, partly filled with SO and partly with a water solution containing non endogenous surface-active molecules and subjected to a sequence of rectangular pulses. The formation of a bulk emulsion was never observed, but droplets were detected at the triple line of contact between the interface and the model solid wall. ^10,11^

Aim of this experimental work was to investigate the role and, for the first time, the interplay of two known major factors influencing SO emulsification, such as eye rotations and albumin as endogenous bio-surfactant.

## 2 Material and Methods

### 2.1 Experimental setup

The eye model has been specifically developed to simulate the real physiological conditions, in terms of temperature and geometry. The model was 3D printed, using Clear Resin 1l (RS-F2-GPCL-04), and it consists of a cylindrical container divided into two identical halves, with an internal cavity that reproduces the human vitreous chamber at real scale (figure 1A). To ensure the model’s sealing, a suitable groove to accommodate an O-ring was prepared on both halves of the model. The vitreous chamber has a realistic geometry and it exhibits the indentation due to the presence of the crystalline lens. ^27–29^ The total volume of the cavity is

**Figure 1:**
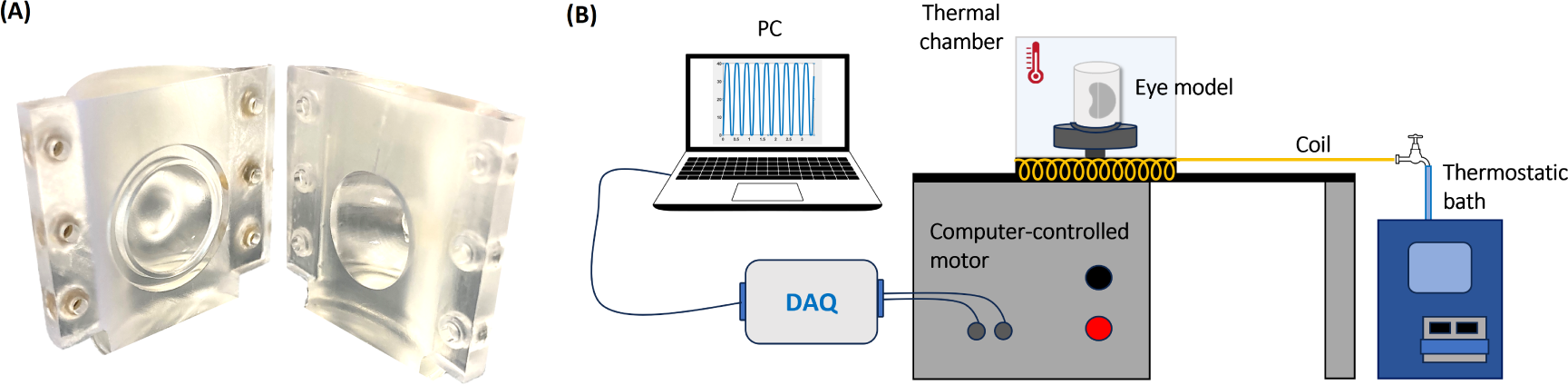
(A) Picture of the unassembled eye model; (B) sketch of the experimental apparatus. *≈* 5.2 ml, in accordance with the recent results of Azhdam et al. ^30^

The eye model was connected to a stepper motor able to reproduce user-defined time law of the angular position. The motor was remotely controlled using a National Instruments^©^ data acquisition system (DAQ) and a Labview^©^ interface to generate the command signal and recording the actual motor shaft position (figure 1B).

To perform the experiments at physiological temperature, the model and the support connected to the motor shaft, were enclosed into a thermostatic chamber, characterised by an insulated base (26*×*26 cm) and a removable Plexiglas covering box. The temperature inside the chamber was maintained at 35*^◦^*C by pumping water from a thermostatic bath (RC 6 CP LAUDA) through a copper serpentine placed within the chamber (figure 1B).

### 2.2 Working fluids and eye model filling protocol

Before each experiment the eye model was carefully cleaned, with standard procedures adopted in surface science laboratories, in order to avoid contamination from surface active molecules and possible impurities. After assembling the eye model, aqueous solution and SO were pumped inside through a hole located on the side of the model, while air flowed out through a second hole, located at top of the model. As during surgery it is never possible to completely fill the vitreous chamber with SO, in the present experiments 20% of aqueous solution was introduced in the eye model. Such solutions containing endogenous molecules were prepared in a Dulbecco phosphate buffered saline (DPBS, Sigma D8662), simulating the in-vivo environment and with different concentrations of bovine serum albumin (Sigma, A2153-50G) equal to 0% (bare buffer), 1% and 5% of the blood concentration (which is *≈* 50 g/l). Albumin was chosen as this protein has a surface-active behavior on the water-oil interface, comparable to that of the mixture of albumin/*γ*-globulins, ^7^ indicating that albumin has a greater influence on the interfacial behavior of molecules present in the serum.

The filling phase required special attention, since we wished to avoid the formation of oil droplets before starting the experiments. Aqueous solution was injected first and then we carefully checked that the inner model’s wall was homogeneously wet. The remaining empty cavity was then very slowly filled with the commercially available SO RS-OIL 1000 cSt (Alchimia srl).

Specific experiments were carried out to evaluate the effect of SO injection (without setting the eye model into motion) on the possible formation of droplets (see table 1).

**Table 1:**
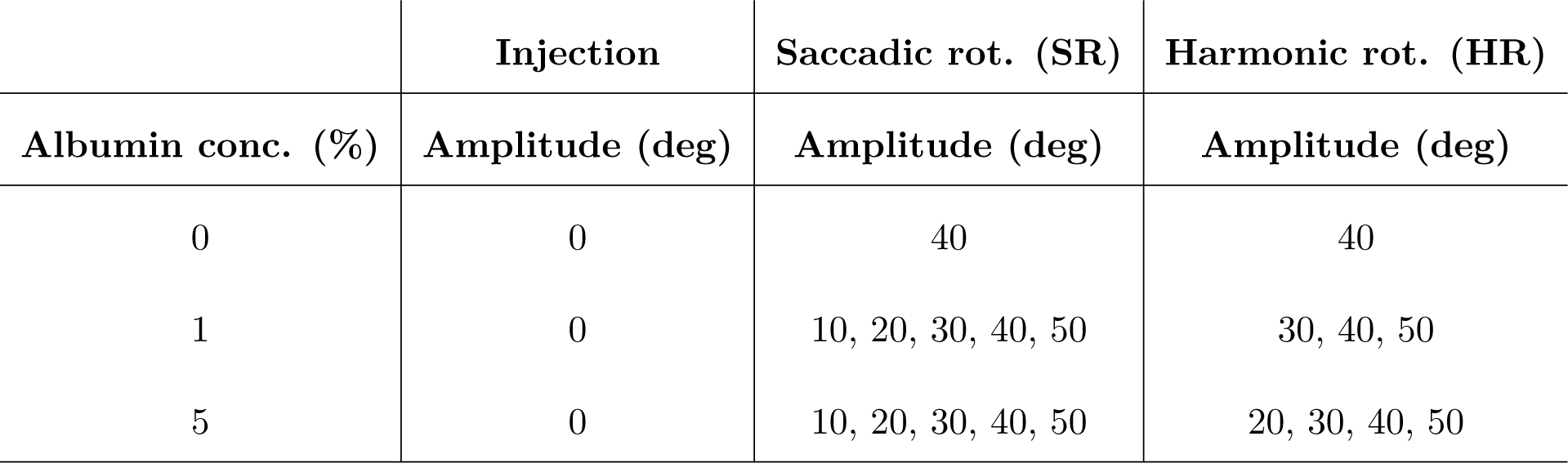
List of the performed experiments.

### 2.3 Simulation of eye movements

Two time laws for simulating eye rotations were considered: harmonic rotations (HR) and a sequence of saccadic rotations (SR) in opposite directions.

A harmonic law is the simplest way to represent a sequence of saccadic eye movements in both directions, with prescribed amplitude and duration. This first set of experiments was run imposing a frequency of 5 Hz and varying the amplitude from 20*^◦^* to 50*^◦^*. Each experiment had a total duration of 1 hour.

A single saccadic rotation is very well represented by the following equation, which was proposed by Repetto et al., ^20^

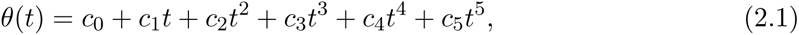

where *θ* is the rotation angle with respect to a given direction and *t* is the time. The 6 coefficients in the above expression can be determined by using metrics of the saccadic rotation, as measured by Becker. ^31^ In particular, he reported that saccade duration *D* linearly increases with the amplitude *A*, according to the relationship

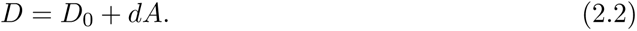

where *A* is measured in degs, *d* assumes the value of 0.0025 s/deg and *D*_0_ is *≈* 0.025.

A periodic signal consisting of a sequence of saccadic rotations in opposite directions has been constructed on the basis of equation (2.1). Specifically, the signal is periodic and a single period consists of a saccade in the clockwise direction, a period of rest of duration *D/*2, a saccade in the counter-clockwise direction and a final additional resting period, again of duration *D/*2. Thus, the period of the signal is equal to 3*D* (see Figure S1 in the Supplementary Material). Since, according to equation (2.2), saccade amplitude and duration are related to each other, once the saccade amplitude *A* is chosen the angular frequency 2*π/*(3*D*) can be calculated. Again the total duration of the experiment was set to 1 hour.

In order to assess the repeatability of the experiments, all tests have been conducted twice, obtaining very similar results, in terms of the observed emulsion. All performed experiments are summarised in table 1.

### 2.4 Microscope image acquisition

At the end of each experiment, the eye model was removed from its support and, 24 hours later, the formed emulsion was extracted and characterised with a microscope quantitative analysis. A DVM6-M optical microscope (Leica, Hamburg, Germany), with a CMOS sensor (3664*×*2748 pixel) and equipped with PlanAPO FOV 3.60 objective, which guarantees a maximum resolution of 2366 lp/mm, was used to determine droplet distribution and size.

Images of the spherical cavity’s content were first captured at 200*×* magnification to locate the emulsion in the vitreous chamber. Once identified, samples were taken and analysed on microscope slides.

Coalescence tests were carried out to determine the type of emulsion formed (water-in-oil W/O vs oil-in-water O/W). A droplet containing the emulsion was placed on a sterile lab slide in contact with a droplet of buffer or SO (see Figure S2 in the Supplementary Material) and the behavior of the system was observed.

For each experiment we performed a droplet size distribution analysis. Specifically, a small amount of the emulsion (about 0.4 ml) was diluted with the continuous phase in order to obtain a good spacing between the droplets, and was then placed between two sterile microscope slides, kept at 0.2 mm distance by a spacer. The volume of the collected sample was equal to *≈* 7.5% of the total volume of the vitreous chamber model and diluted with an amount of aqueous solution variable from case to case. Although the total volume sampled could not allow for an accurate quantification of the total amount of emulsion, the adopted method is reliable and reproducible in terms of droplet size distribution, which is one of the findings we focused on.

Using the Leica Application Suite X proprietary software, micro-photos of the emulsions were taken, with different magnifications (700*×*, 900*×*, 1200*×* and 1800*×*) with resulting spatial resolution in the acquired image ranging from 0.2 to 0.7 *µ*m. These spatial resolutions are comparable with the physical limits of the optical microscope. Owing to these resolution constraints, we limited our analysis to drops with diameters *≥* 1 *µ*m. For each sample, about 15 images were acquired, which was sufficient to cover the entire sample.

The images were then analysed through the same software, to determine droplets size distribution. A specific tool of this software allows the operator to identify all drops within an image and it returns the droplets diameter. Since SO drops reflected the microscope light, droplet boundaries could vary from 1 to 2 *µ*m in thickness. For this analysis analysis droplets external diameter was considered.

The size of the observed droplets was analysed in term of size frequency distribution using a bin size of 5 *µ*m, which is significantly larger than the experimental error (which is of the order of 1 *µ*m).

For each test, we also computed the surface occupied by the droplets over the total area of the acquired images, *S*. This quantity was computed as

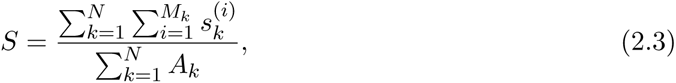

where *N* is the number of considered images, *M_k_* is the number of droplets in the *k*-th image, *A_k_* is the area of the *k*-th image and 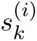 is the area of the *i*-th droplet in the *k*-th image. This dimensionless number *S* allowed us to compare images obtained with different magnifications in terms of area occupied by the droplets.

The performed statistical analysis was based on the hypothesis that the drops observed on the focal plane of interest were sufficiently representative of the emulsion; this aspect was confirmed by the repeatability of the experiments.

## 3 Results

No emulsions were detected in the absence of endogenous proteins in aqueous solution; whereas, emulsified SO droplets formed in all the tests with aqueous solution containing albumin. Remarkably, this happened for all rotations imposed to the eye model and all albumin concentrations (1% and 5% of blood serum concentration). In general, we predominantly observed the formation of SO droplets in the region of the geometric indentation, which reproduces the crystalline lens. This is the site where the flow field induced by eye rotations has a complicated three-dimensional structure. ^14,15^

In experiments without eye motion, a very small amount of SO droplets was detected in the case of the highest concentration of albumin (5%) (see Figure S3 of the Supplementary Material), confirming that the emulsion observed in the experiments described in the following can almost entirely be attributed to the motion of the eye model.

Interestingly, coalescence tests confirmed that all the emulsions were of SO in aqueous solution type (O/W) as we invariably observed that, a droplet of the emulsion coalesces with an adjacent drop of buffer, leading to a droplet distancing; this did not happen when SO was added (see Figure S2 in the Supplementary Material).

We kept the formed emulsions in static conditions at room temperature for a week and did not observe any appreciable change in drop number and size, which shows that the emulsions are stable over long time scales.

Figure 2 includes representative images acquired with the optical microscope, showing the presence of different drops size, as well as sub-micrometric droplets, below the software resolution of about 1 *µ*m for the magnification used. Thus, droplet sizes *≤* 1 *µ*m are not included in the size distribution discussed below.

**Figure 2:**
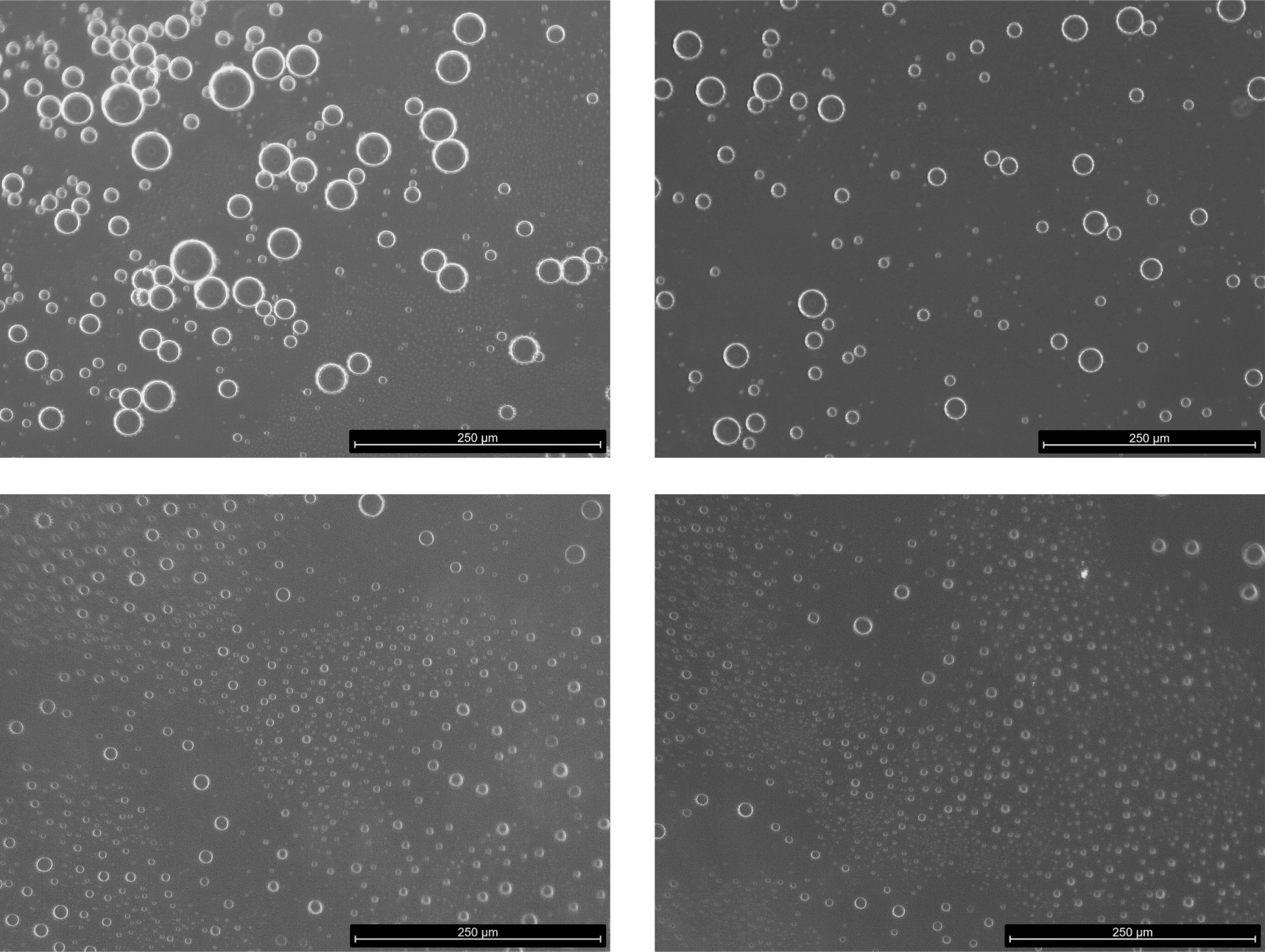
Examples of microscope images of the emulsion formed dilute with the continuous phase for droplet distribution analysis. HR experiemnt with albumin at 1% and *A* = 40*^◦^*.

Figure 3 shows droplet distributions (in %) at the two investigated albumin concentrations and the two types of imposed rotation, SR (Figure 3 A,B) and HR (Figure 3 C,D). Droplet size distributions obtained with SR (Figure 3 A,B) qualitatively resemble a Poisson distribution. The majority of drops has a very small diameter and the most frequent diameter class is 6 to 10 *µ*m. No particular differences were observed between the two albumin tested concentrations (see also Figure S4 in the Supplementary Material). The distributions seem to suggest that increasing the amplitude of eye rotations leads to an increase in the number of small droplets, though differences between different rotation amplitudes are subtle and not all curves reported in the figures are ordered accordingly. The dependency on the eye rotations amplitudes is clearer for HR when SO droplets get smaller as the amplitude increases. This is because for HR the amplitude is changed keeping the frequency fixed, which is not the case for the SR, as explained in §2.3. For rotations amplitudes of 40*^◦^* and 50*^◦^* we didn’t manage to resolve the left, growing branch of the curves, which suggests that a large number of formed droplets were too small to be detected, likely due to the more intense motion imposed in the case of HR than that corresponding to SR (see also Figures S4 and S5 in the Supplementary material). In the case of large amplitude oscillations, the motion is in fact too intense to represent physiological conditions. However, HR experiments have the value to clearly demonstrate that the intensity of eye movements has a key role in the process of droplet generation and that the higher the mechanical energy put into the system, the smaller the average droplet size. In HR experiments also albumin concentration plays a more recognisable role, as shown by the black curve corresponding to 30*^◦^* amplitude rotations (Figure 3), strongly displaced to the left moving from concentration of 1% to 5%, and almost collapsing on the other curves, corresponding to rotations of larger amplitude.

**Figure 3:**
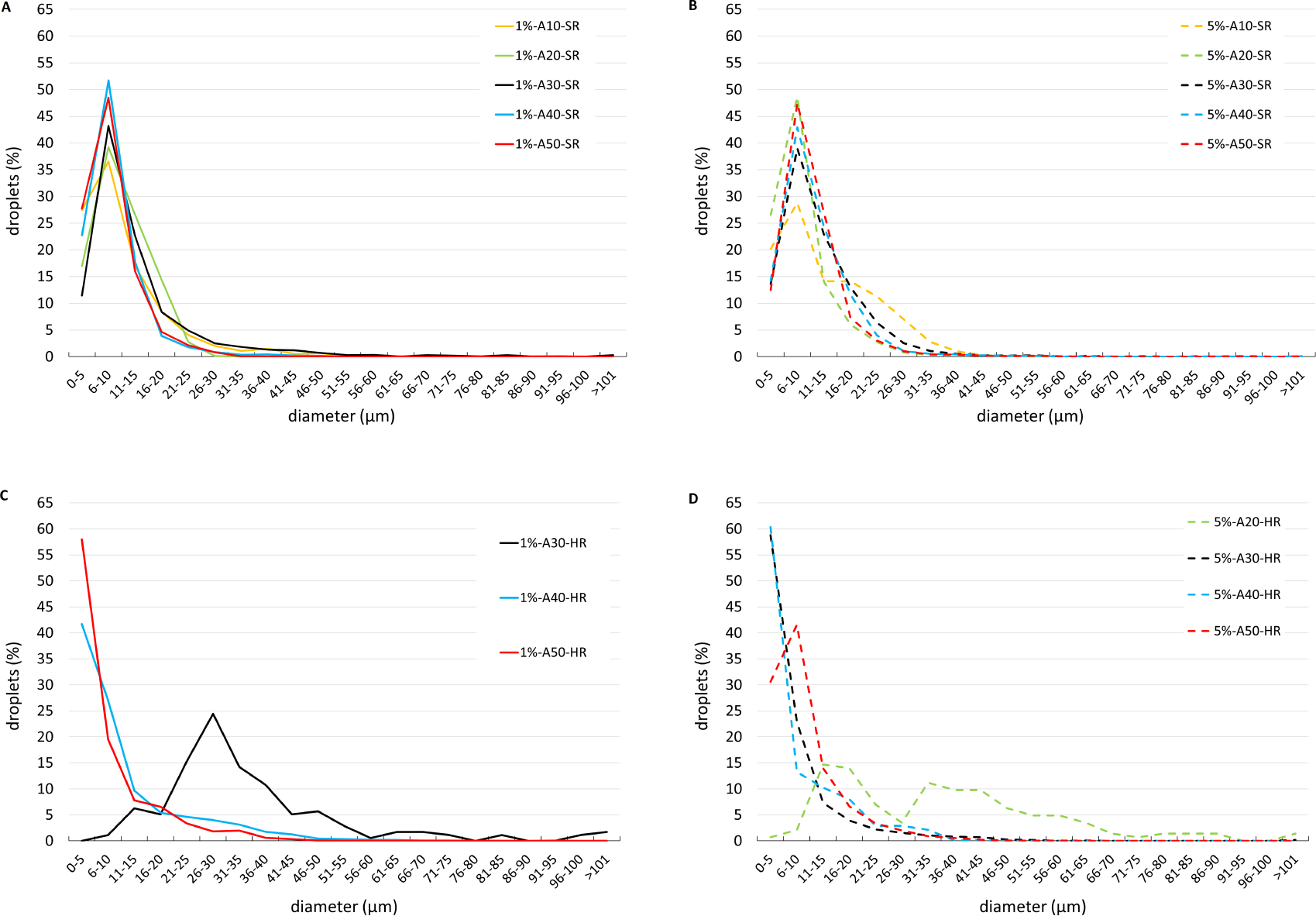
Comparison of droplets size distributions (in %) of emulsions obtained through SR experiments (A,B) and HR experiments (C, D), for all the investigated rotation amplitudes. Solutions containing 1% of albumin (A,C) and 5% of albumin (B,D). Different color bars correspond to different imposed amplitudes.

The results shown in the Figure 4 confirmed the finding discussed above. In particular, the dimensionless parameter *S* and the droplet density grew with the rotation amplitude in the case of HR but not of SR. The effect of albumin concentration was also more evident in the case of HR (Figure 4).

**Figure 4:**
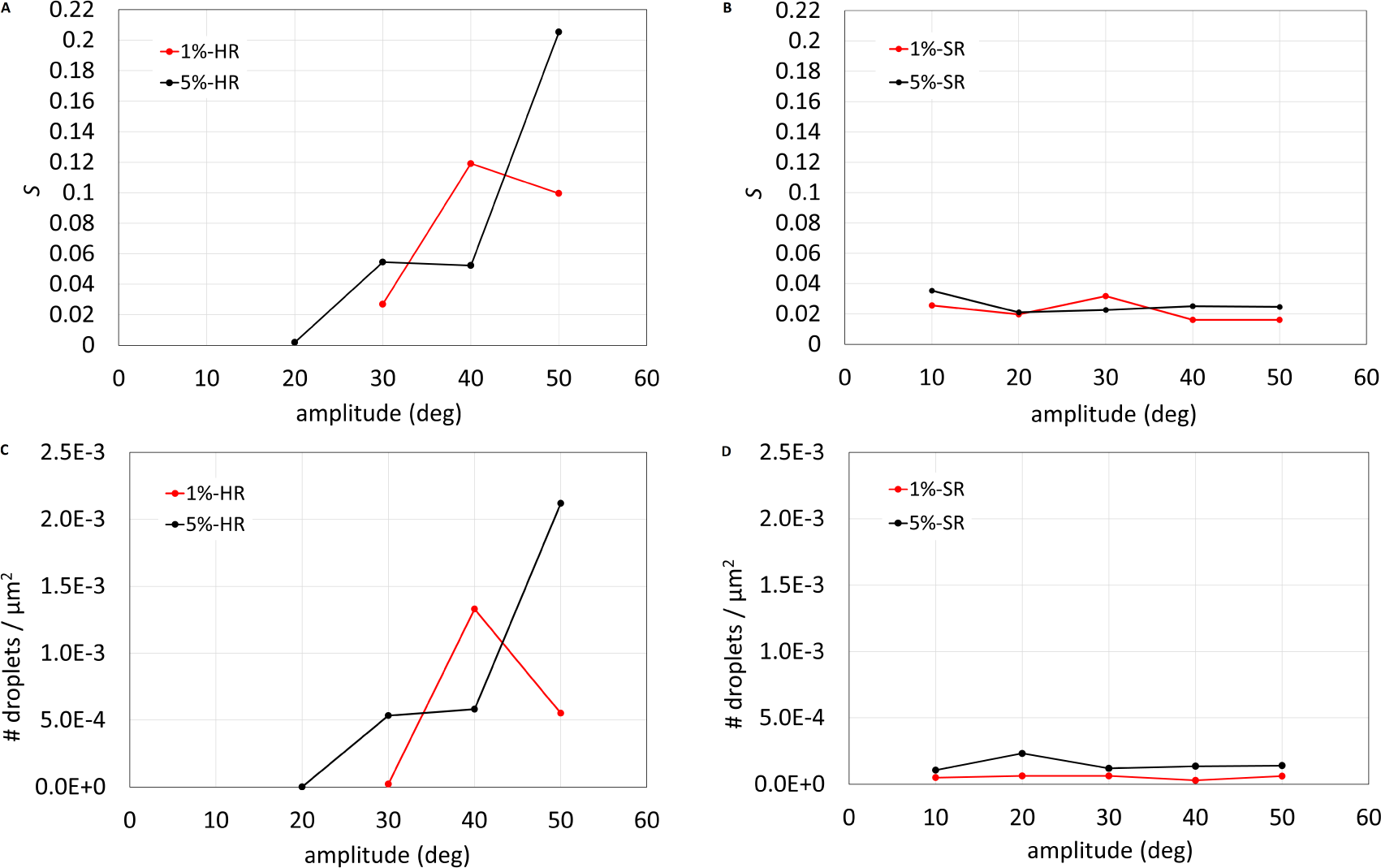
(A,C) HR experiments, (B,D) SR experiments. (A,B) Surface area occupied by droplets with respect to the area of the acquired images (*S*), as a function of the imposed rotation amplitude. (C,D) Total number of droplets per unit area as a function of the imposed rotation amplitude. Different colors correspond to different albumin concentration.

## 4 Discussion

In this study we investigated the formation of SO emulsion in a vitreous chamber model, focusing on the influence of eye rotations, albumin and combination of both. Although similar experiments have been previously conducted ^10,11^ our study presents some key differences. First, our model is not spherical but has a realistic geometry of the human vitreous chamber. Analytical ^14,15^ and experimental^21^ works have shown that even a weak departure of the vitreous chamber shape from spherical has a very significant effect on vitreous motion, causing the formation of fluid circulations in the anterior part of the vitreous chamber, due to the change in curvature of the wall produced by the crystalline lens. Second, we kept the model at body temperature during all the experiments, which possibly has a major effect on the formation of an emulsion. In fact, the chemico-physical properties of SO-aqueous interface are strongly dependent on temperature. ^32^ Third, we use aqueous solutions containing two different concentrations of albumin, due to the demonstrated major role on interfacial properties. ^7^ Finally, realistic saccadic rotations have been imposed on the eye model.

The eye model was set in motion following both simple harmonic time laws (HR) and also realistic sequence of saccadic rotations (SR). All experiments lasted an hour, thus consisting of a very long series of successive eye movements.

In the absence of albumin we never found formation of SO droplets in the aqueous solution, which confirms its key role in the emulsification process. We note that, though albumin is not normally present in the aqueous humour, ^33^ its presence in the vitreous chamber after PPV is almost certainly unavoidable, owing to possible bleeding and the postoperative inflammatory processes. ^1^

In all experiments we followed the same SO injection procedure and we verified that the injection phase alone did not produce bulk emulsification but a very small amount of droplets in the vitreous chamber. This confirmed that the majority of observed drops in the experiments can be attributed to the mechanical energy generated by eye movements. We note that we took care to avoid SO drop formation during the filling phase, which was not designed to faithfully reproduce the SO injection during surgery. Thus, our results do not provide indications as to whether SO droplets can form during injection in the real surgery.

In all experiments in which the eye model was set in motion, we found by coalescence experiments the formation of SO oil-in-water (O/W) emulsion, in agreement with the Bancroft rule ^34^ (and not as hypothesised elsewhere). ^35^

Different research groups carried out similar experiments and did not observe the formation of bulk emulsions, but the generation of droplets only in the vicinity of the triple line of contact of the interface with the wall. ^10,11,36^ The authors provide sound physical explanation of how the movement of the triple line could cause the formation of SO droplets. The discrepancies with our results are likely to be attributed to the use of a model of the vitreous chamber with realistic geometry and kept close to body temperature. Finally, instead of non-endogenous, surface-active molecules, ^10,11^ we employed an aqueous solution containing albumin, which has previously been shown to significantly affect the properties of the SO aqueous solution interface. ^7^ All the above ingredients make the formation of emulsions more likely.

The characterisation of the size distribution of the observed SO droplets showed that drops were of a very small size, with the class comprising the majority of drops typically being below 10 *µ*m. Furthermore, areas composed of submicrometric droplets were observed. This is in line with previous studies demonstrating that small SO droplets (*<* 2 *µ*m) represent the major component of SO emulsification *in vivo* ^37,38^ and the demonstrated positive correlation between SO droplets *<* 2 *µ*m and larger ones (7-30 *µ*m), ^37^ reinforces the translational value of our research in terms of characterisation of SO emulsion. From a clinical point of view, emulsified SO microdropltes may be particularly dangerous as they can be phagocytosed by macrophages, resident microglial and RPE cells, thus, promoting the SO-related intraocular inflammation and, in turn, further SO emulsification. ^3^ In addition, the detection of emulsified SO droplets within ocular structures of both anterior and posterior segment, support not only their ability to penetrate tissues, but also their relevant contribution to complications associated with SO use. ^37,39,40^ For instance, emulsification, inflammation and mechanical obstruction of the trabecular meshwork, appear to have a fundamental role in SO-related IOP elevation and secondary glaucoma. ^41^ The migration of emulsified SO droplets and the consequent contact with corneal endothelium may have a crucial role in SO-related corneal changes. ^42^ Epiretinal, intraretinal and subretinal emulsified SO droplets have been detected as hyperreflective dots on spectral domain-optical coherence tomography (SD-OCT) ^43,44^ and the SO-related thinning of inner retinal layers is one of the mechanisms potentially involved in SO-related vision loss. ^2^

In the experiments in which the eye model was subjected to HR of large frequency and amplitude, SO drops became too small to be counted and measured with the optical techniques employed in this work. The general indication, is that the higher the amount of mechanical energy put into the system and the higher albumin concentration in the aqueous phase, the small the size of droplets formed. We note that we tested HR and SR but also other ocular movements are likely to play a role in the formation of an emulsion, most notably REMs. ^45^

With regard to albumin, differences between concentrations of 1 or 5% of that in serum blood are subtle, especially in the case of SR. This suggests that even a very small amount of proteins in the aqueous solution is sufficient to induce changed in the rheological properties of the interface strong enough to lead to emulsion formation. Clinically, these results may highlight the relevance of surgery- and patient-specific findings in the development of SO emulsification. Indeed, the minimisation of intraoperative bleeding and an optimised control of inflammation after PPV might minimise the potential presence of albumin (as well as other blood serum proteins) into the vitreous chamber and, thus, contribute to lower the rate of postoperative SO emulsification. Finally, the verified stability over time of SO emulsion may support its potential to induce long-term detrimental effects.

## Acknowledgments

The authors acknowledge Alchimia srl (Italy), for supplying SO samples.

