## Supplementary figures for "The role of eye movements in the process of silicone oil emulsification after vitreoretinal surgery"

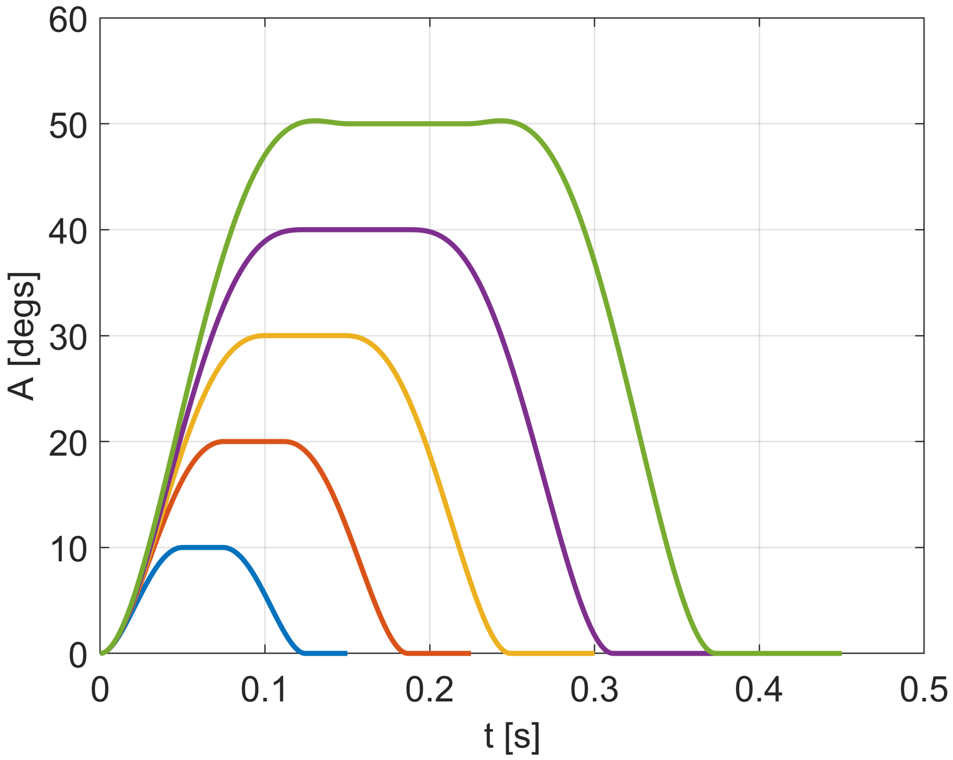


Figure S. 1: Single period of the saccade signal used for the SR experiments with $A$ imposed between 10° and 50°. The period of the signal is equal to $3D$ and consists in a saccade in the clockwise direction, a period of rest of duration $D/2$, a saccade in the counter-clockwise direction and a final additional resting period of duration $D/2$.

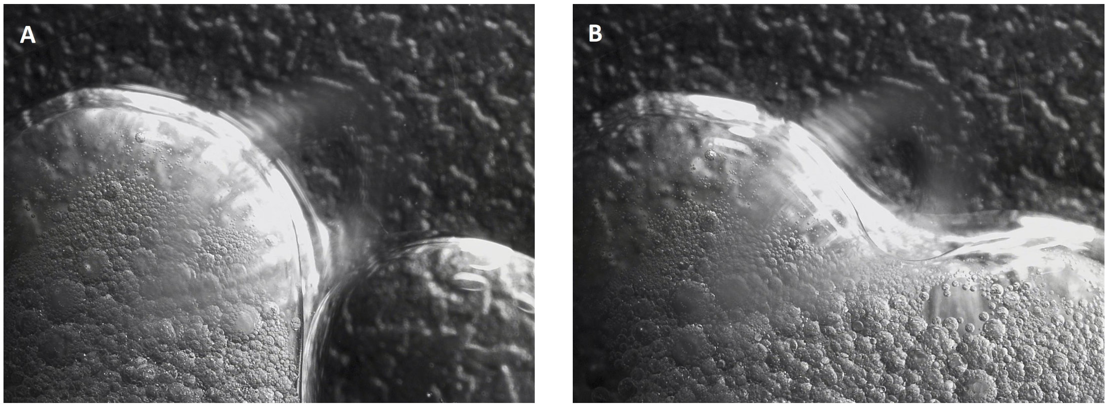


Figure S. 2: Consecutive microscope acquired frames of the coalescence test. (A) DPBS droplet (right) positioned close to the droplet of emulsion (left); (B) Occurred colescence between the two droplets. This allows to consider the emulsion as oil-in-water (O/W) type. To verify this result the same test has been performed positioning an oil droplet close to a droplet of emulsion. In this case, these two phases di not coalesce, confirming that the continuous phase of the emulsion is the DPBS solution.

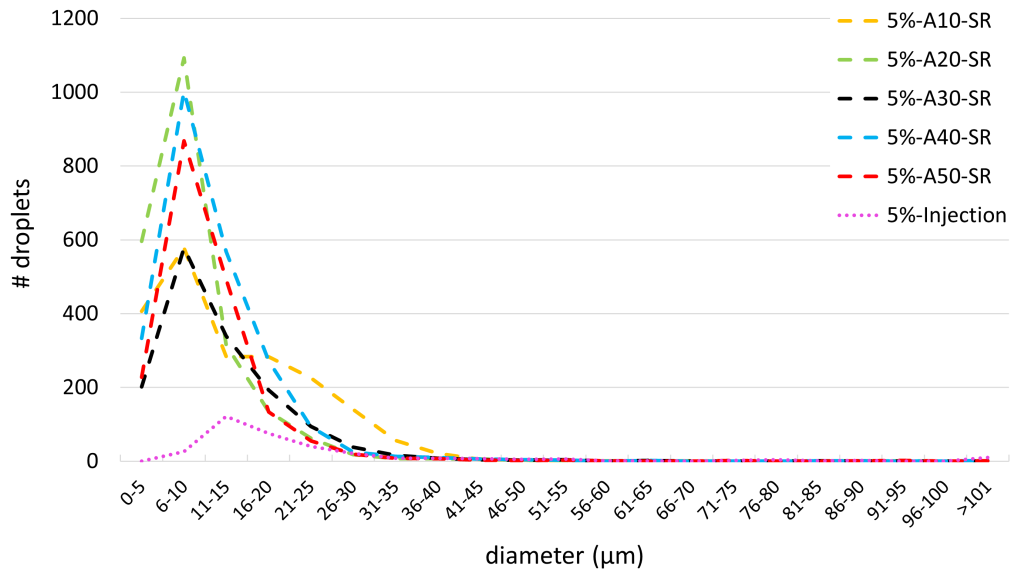


Figure S. 3: Comparison of droplets size distributions (in number of droplets) of emulsions obtained through SR experiments and with 5% of albumin concentration for all the investigated rotation amplitudes. Different color bars correspond to different imposed amplitudes. In the figure the droplets size distribution of drops observed after the injection phase is also reported.

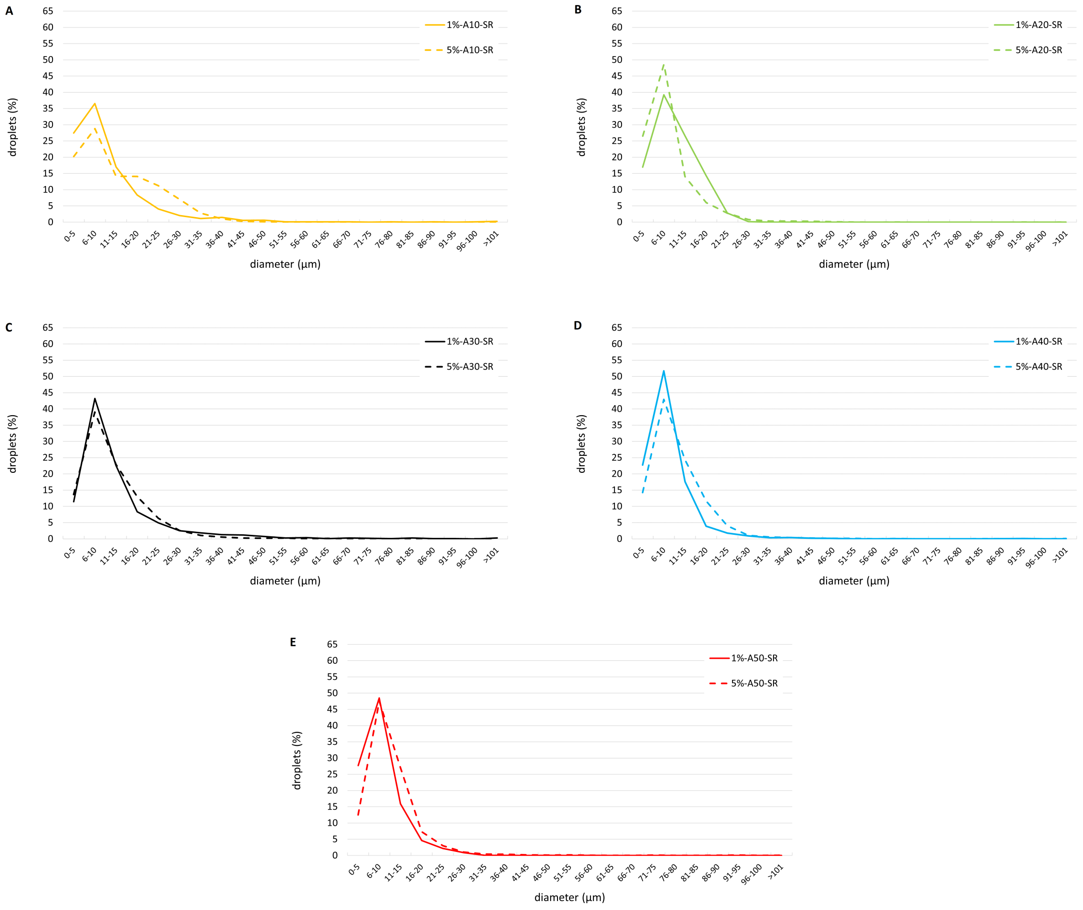


Figure S. 4: Comparison of droplets dimension distribution (in %) related to SR experiments with albumin concentration at 1% (continuous lines) and at 5% (dashed lines). Each panel reports the results obtained by imposing the same saccadic amplitude: (A) $A$=10°, (B) $A$=20°, (C) $A$=30°, (D) $A$=40°, (E) $A$=50°. Results obtained with the two albumin concentrations resulted very similar for all the imposed rotation amplitudes.

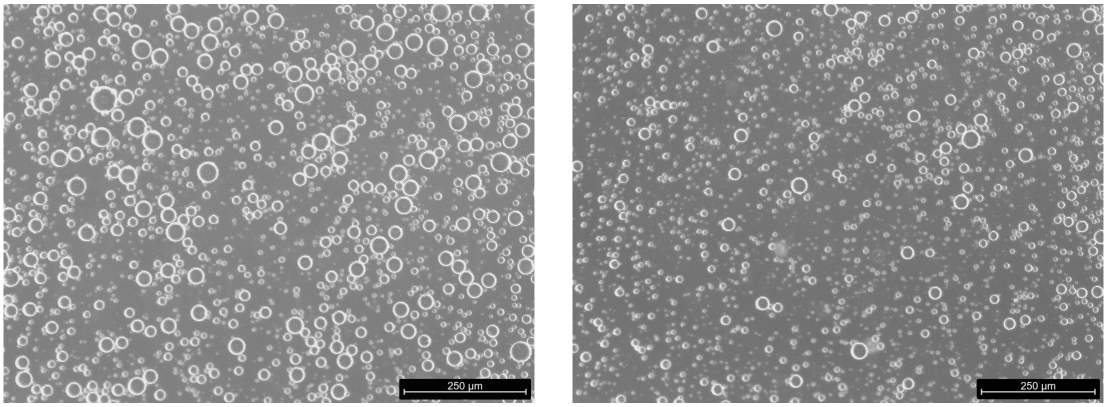


Figure S. 5: Example of microscope images of emulsions obtained through HR experiments with $A$=50°. A significantly larger area occupied by SO droplets for each image can be observed with respect the images acquired with lower HR amplitudes for the same albumin concentration.
